# The transition state for coupled folding and binding of a disordered DNA binding domain resembles the unbound state

**DOI:** 10.1101/2024.02.12.579954

**Authors:** Mikhail Kuravsky, Conor Kelly, Christina Redfield, Sarah L Shammas

**Affiliations:** Department of Biochemistry, University of Oxford, Oxford, OX1 3QU, UK

## Abstract

The basic zippers (bZIPs) are one of two large eukaryotic families of transcription factors whose DNA binding domains are disordered in isolation but fold into stable α-helices upon target DNA binding. Here we systematically disrupt pre-existing helical propensity within the DNA binding region of the homodimeric bZIP domain of cAMP-response element binding protein (CREB) using Ala-Gly scanning and examine the impact on target binding kinetics. We find that the secondary structure of the transition state strongly resembles that of the unbound state. The closest residue to the dimerisation domain that has been examined is largely folded within both unbound and transition states; dimerisation apparently propagates additional helical propensity into the basic region. The results are consistent with electrostatically-enhanced DNA binding, followed by rapid folding from the folded zipper outwards. Interestingly, despite taking the exact experimental approach suggested for testing it, we find no evidence for disorder-mediated rate enhancement predicted by fly-casting theory.

## INTRODUCTION

Eukaryotic transcription factors frequently contain intrinsically disordered regions that fold upon binding to target DNA sequences; the AT-hooks, bZIP and bHLH domains (1). In the absence of DNA, the basic DNA binding domains are dynamic but exhibit varying levels of pre-existing helical content (2–4). The existence of structures in the unbound state that resemble final folded structures accelerate binding processes if these are the ones selected for binding in a conformational selection mechanism (5). In contrast if the domain only folds after binding in an induced fit mechanism then pre-existing structures may have little impact on association rates (6–8). Though not yet experimentally verified, protein disorder has even been speculated to be advantageous to association rates through a mechanism known as fly-casting (9). This mechanism involves region(s) of the extended disordered protein/region first binding to their partner with the rest of the protein being subsequently “reeled in” and folding. Thus, the relevance of pre-existing structure in regions that fold upon binding could be intimately linked with folding mechanisms.

A protein’s structure, or fold, is encoded in its amino acid sequence. But how are these folds achieved? Proteins have open to them an immense conformational space, yet natural proteins converge to their native states in sub-second timescales. A powerful experimental technique to probe protein folding mechanisms has undoubtedly been the Φ-value analysis (10). This mutation-based method was developed to describe the folding transition state. Conservative mutations are made to the amino acid sequence, and the effect on folding/unfolding rates is compared to the overall free energy change. The resulting Φ - value, which typically lies in the range 0-1, describes the extent of eventual (native) contacts that the residue has acquired by the transition state (10). Over the last ten years this method has now also been applied to investigate the coupled folding and binding of a handful of intrinsically disordered proteins/regions (IDPs/IDRs) with their partner proteins (6, 7, 11–21), however the transition state for folding of a protein-DNA interaction has not yet been reported.

One protein containing a disordered DNA binding domain is the cAMP-response element binding protein (CREB). CREB is an ubiquitously expressed transcription factor regulating expression of a wide range of genes (22–24) through binding to cAMP-response element sites (CRE, consensus 5’-TGACGTCA-3’) in the proximal promoters of its targets (25, 26). Phosphorylation at S133 within the Kinase Inducible Domain (KID) of CREB promotes interactions with its coactivator, CREB-binding protein (CBP) (27, 28), enabling transcription by recruiting the core transcription machinery (29). CREB acts as a hub in a signalling network controlling proliferation (30), differentiation (31–33) and the survival of cells (34, 35). Transcriptional activity of CREB plays a central role in a number of processes, including immune response (36), reproduction (37, 38), long-term memory (39) and circadian rhythms (40).

DNA binding by CREB is mediated by a C-terminal basic leucine zipper (bZIP) domain (41), part of a eukaryotic superfamily of transcription factors that has 56 human members (42). A typical bZIP binds to the DNA as a homo- or heterodimer. Each of the subunits consists of an N-terminal basic region (BR) and a C-terminal leucine zipper (LZ) (Figure 1A, B). The LZ promotes dimerisation and is built of two amphipathic α-helices wrapped around each other in a coiled-coil structure. The BR mediates DNA recognition and is enriched in positively charged amino acids (43). As shown in a number of X-ray structures, a bZIP assumes a chopstick-like structure of two uninterrupted α-helices grasping the DNA by the major groove (44–51). Although the bZIP domains have been studied for more than 30 years, there are few details on the mechanism of their binding to DNA. Here we perform mutational scanning of the BR of homodimeric CREB bZIP to probe its binding mechanism and reveal the extent of helical formation within the transition state for binding.

**Figure 1.**
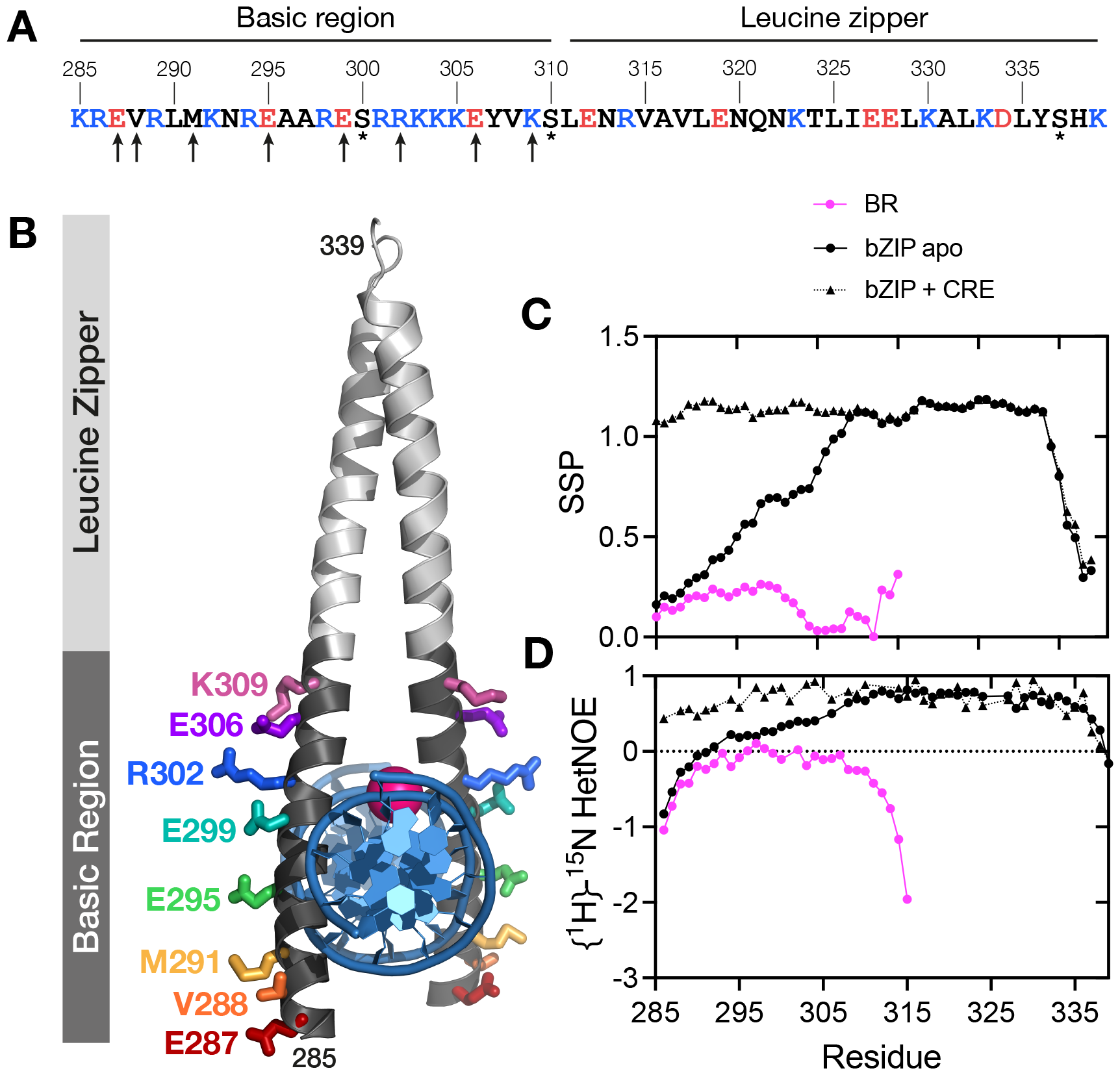
Structural description of CREB bZIP. A. The sequence of the CREB _285_bZIP construct used in this study. Positively charged amino acids are shown in blue, negatively charged amino acids are shown in red. All cysteines were replaced with serines (asterisks). Eight solvent-exposed residues subjected to alanine and glycine substitutions are marked with arrows. B. Structural positions of residues subjected to mutagenesis within CREB _285_bZIP-DNA complex. The Mg^2+^ ion is shown as magenta sphere. Derived from 1DH3. Calculated Secondary Structural Propensity (C) and {^1^H}-^15^N heteronuclear NOE (D) of CREB _285_BR (pink circles), CREB _285_bZIP (black circles) and CREB _285_bZIP with CRE DNA (black triangles).

## MATERIAL AND METHODS

### Cloning

A synthetic gene for the bZIP domain of human CREB (_285_bZIP) corresponding to residues 285-339 of CREB-A (Uniprot P16220) was fused with the B1 domain of Streptococcal protein G (GB1) at the N-terminus and cloned into a pRSET-A vector. A tobacco etch virus (TEV) protease site was inserted between the GB1 and the bZIP sequence. The C300, C310 and C337 residues of _285_bZIP were mutated to serine as described in (52) to improve protein solubility without affecting its DNA binding properties. Plasmids encoding mutant versions of _285_bZIP were produced by point mutagenesis and verified by DNA sequencing. The monomeric basic region of CREB bZIP corresponding to residues 285-315 of CREB-A was generated by insertion of a stop codon. In the following, the numbering of amino acids is consistent with that of the longest CREB isoform (CREB-A).

### Protein expression and purification

Plasmids for the wild-type and mutant bZIP constructs were transformed into *Escherichia coli* BL21(DE3)pLysS. Cells were cultured at 37 °C in 2xYT medium. After the OD_600_ reached 0.6-0.8, recombinant protein expression was induced by 1 mM IPTG, and culturing was continued for another 4 h. *E. coli* pellets were disrupted by sonication in 100 mM Tris-HCl pH 8.5, 50 mM NaCl (binding buffer). The lysate was cleared by centrifugation and loaded on a Ni-NTA agarose column (HisTrap HP; GE Healthcare). After rigorous washing with 100 mM Tris-HCl pH 8.5, 2 M NaCl to remove bacterial DNA, protein was eluted with a 0-2 M gradient of imidazole in the binding buffer. His6-GB1-tag was subsequently cleaved by TEV protease (3 mM) and separated from bZIP on a cation exchange column (HiTrap SP; GE Healthcare) using a 0–2 M gradient of sodium chloride. Finally, proteins were buffer exchanged into 10 mM MES pH 6.5, 150 mM NaCl, 10 mM MgCl_2_, 0.05% Tween-20 (biophysical buffer). Protein purity was confirmed by SDS-PAGE (Supplementary Figure 1) and electrospray ionization mass spectrometry (ESI-MS). The absence of nucleic acid contamination was assessed by measuring the 260/280 ratio to be 0.42. All protein constructs contain an additional glycine residue at the N-terminus as a remnant from TEV cleavage.

### Expression of isotopically labelled CREB

Isotope labelling used a modified version of the protocol by (53). In short, cells were grown in 1 L of 2xTY at 37 °C, shaken at 180 rpm. Once OD_600_ ∼0.7 was reached, cells were pelleted at 5000 g for 15 min at room temperature, washed with 1x M9 minimal media and resuspended in 250 mL of 1x M9 minimal media containing FeCl_3_, MgSO_4_ and CaCl_2_ (1 mM respectively), supplemented with 4 g/L U-^13^C D-glucose (Cambridge Isotopes CLM-1396) and/or 1 g/L ^15^N NH_4_Cl (Cambridge Isotopes NLM-467). Cells were then incubated for 45 min at 37 °C shaken at 180 rpm, induced with 1 mM IPTG and cultured for a further 4 hours. Protein purification was then carried out as described above.

### Protein concentrations

The molar extinction coefficient of CREB _285_bZIP in biophysical buffer was determined as 2560 M^−1^cm^−1^by Gill and von Hippel method (54).

### Preparation of double-stranded DNA

The binding of CREB to a consensus-CRE-containing DNA sequence was examined using self-annealing CRE oligonucleotides (5’-CC**TGACGTCA**GCCCCC**TGACGTCA**GG-3’) prepared according to (55) and labelled with Alexa Fluor®488 at the 5’-terminus (Alexa Fluor®488-CREh). Dissociation kinetics were studied using a 14 bp double stranded competitor CRE DNA (5’-CC**TGACGTCA**TCCG-3’). Out-competition using identical (non-labelled) sequences results in additional phases in stopped-flow experiments that are also present without CREB. Finally, NMR studies utilized a symmetric CRE-containing sequence (5’-AGA**TGACGTCA**TCT-3’). Annealing was performed by mixing the sense and antisense strands at 100 µM in water, heating to 95 °C and subsequently cooling down to 4 °C at a rate of 0.1 °C/min. All DNA was purchased from Life Technologies.

### Equilibrium dissociation constant for homodimerisation

Homodimer formation was monitored by following the change in intrinsic tyrosine fluorescence on an SX20 stopped-flow spectrophotometer (Applied Photophysics, Leatherhead, UK). CREB 285-339 (_285_bZIP) in 8 M urea was mixed rapidly in a 1:10 volume ratio with buffer solutions of various (lower) urea concentrations, to achieve protein and urea concentrations outlined in the text. An excitation wavelength of 278 nm was used with a 305 nm cut-off filter, and the temperature was maintained at 25.0 °C. At least 50 traces were averaged for a typical measurement.

Kinetic refolding traces were fit using proFit (QuantumSoft, Tomsk, Russia) software to

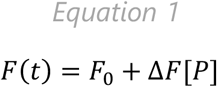

where the concentration of monomeric CREB, [P], is given by 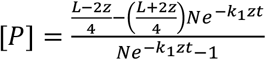

*L* is the homodimerisation equilibrium constant, *k*_1_ is the folding rate constant, 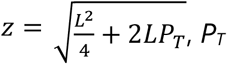 is the total CREB concentration, *N* is 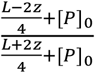, and due to the conditions of our refolding experiment *P*_0_ = *P*_T_. The equation was derived as shown in the Supplementary Material.

The value of *L* in the absence of urea, and the equilibrium m-value for homodimerisation, were extracted as the y-intercept and gradient of a straight line fit of L against urea concentration. The value of *k*_1_ in the absence of urea, and the kinetic m-value for folding, were extracted as the y-intercept and gradient of a straight line fit of *k*_1_ against urea concentration.

### Circular dichroism

The measurements were carried out for 20 µM protein samples in 10 mM MES pH 6.5, 150 mM NaCl, 10 mM MgCl_2_, 0.05% Tween-20 (biophysical buffer) at 25 °C. Spectra were collected using a Jasco J-815 spectropolarimeter equipped with a 1 mm pathlength cuvette and corrected for buffer contribution. Fractional helicities (FH) were calculated using the method described in (56).

### Nuclear Magnetic Resonance (NMR) Spectroscopy and Secondary Structure Propensity Calculations

NMR experiments were carried out using spectrometers operating at ^1^H frequencies of 600 and 750 MHz. The spectrometers were equipped with Oxford Instruments magnets and Bruker Avance III HD consoles with TCI cryoprobes. Data were acquired using pulse sequences in the Topspin library from Bruker Biospin, or using the BEST-TROSY library (57), with non-uniform sampling for most triple resonance experiments. Data processing was performed using NMRPipe (58), using istHMS for reconstruction of NUS acquired data (59), with visualisation and assignments being performed using CCPN Analysis v2.4.2 (60). Sequential assignments were obtained using 2D and 3D double and triple resonance experiments, including ^1^H-^15^N-HSQC, ^1^H-^15^N-BTROSY, ^1^H-^13^C-HSQC, ^15^N-edited TOCSY-HSQC, ^15^N-edited NOESY-HSQC, HNCO, HN(CA)CO, HNCA, CBCA(CO)NH, HN(CO)CACB, HBHA(CO)NH, HCCH-TOCSY, HCC(CO)NH, and (H)CC(CO)NH. The {^1^H}-^15^N heteronuclear NOE was measured at 25 °C in an interleaved experiment recorded with and without ^1^H saturation for 3.5 s at 600 MHz. NMR samples contained 0.3-1 mM ^15^N or ^13^C/^15^N-labelled CREB constructs. The DNA-bound state was examined using a two-fold excess of unlabelled DNA over CREB dimer. Buffer used for NMR was 95% H_2_O/5% D_2_O with 10 mM Tris-d_11_ (Sigma), 150 mM NaCl, 10 mM MgCl_2_, 1 mM NaN_3_ plus protease inhibitors (Pierce™, Thermo Scientific). Spectra were referenced to DSS. Details of assignments for each specific complex are detailed in BMRB depositions 50880, 50872, 50873. Secondary Structure Propensities were calculated using the ^13^Ca and ^13^CO chemical shifts using the SSP program from the Forman-Kay laboratory (61).

### Kinetic measurements of CRE binding and unbinding

Kinetics experiments were carried out in biophysical buffer at 25 °C using an SX20 stopped-flow spectrometer (Applied Photophysics). Reaction progress was followed by monitoring the change in fluorescence intensity of the labelled DNA using excitation wavelength of 495 nm and a 515 nm long-pass filter. Association measurements were conducted under the pseudo-first order conditions by rapidly mixing 10 nM DNA with 100-200 nM protein in a 1:1 ratio. The averaged traces were fit with a single-exponential decay function to obtain the observed rate constants, and association rate constant (*k*_on_) values were determined from the gradients of linear fits to observed rate constants plotted against protein concentrations. Dissociation measurements were performed by mixing the pre-formed protein-DNA complex (100 and 10 nM, respectively) with a saturating excess of unlabelled competitor CRE DNA (4 µM). For the most severely destabilising mutations, the concentrations of all reaction components were doubled. Dissociation rate constant (*k*_off_) values were obtained by fitting the averaged traces with a single-exponential decay function. Equilibrium dissociation constants (*K*_D_) for CRE DNA were estimated as the ratios of dissociation to association rates.

### Φ-value analysis

Φ values were calculated according to (11, 12):

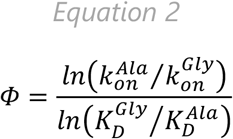

where 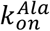 and 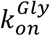 are association rate constants for alanine and glycine mutants, respectively, whilst 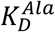 and 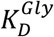 are equilibrium dissociation constants for alanine and glycine mutants, respectively. Φ value for R302 was calculated using dissociation constants obtained from equilibrium titrations; Φ values for the rest of the substitutions were calculated using kinetic estimates of the 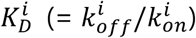

### Equilibrium titrations

Equilibrium binding of CREB bZIP to CRE DNA was measured by following steady-state fluorescence anisotropy of labelled DNA probes. Briefly, the protein was titrated to 5 nM AlexaFluor®488-CREh oligonucleotides in biophysical buffer. Samples were equilibrated at 25 °C for 1 min and assayed using a PerkinElmer LS-55 spectrofluorometer equipped with a 3 mm square cell. Excitation and emission wavelengths were set at 495 and 519 nm, respectively.

The data were analysed according to a model assuming that CREB binds to DNA as a preformed dimer and accounting for a secondary binding event that was observed at high protein concentrations:

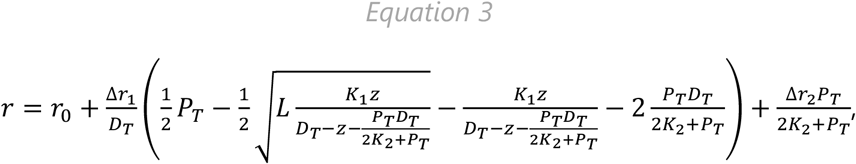

where 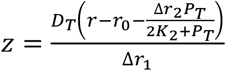, *r*_0_ is the anisotropy of free DNA, Δ*r*_1_ and Δ*r*_2_ are the signal changes upon the primary and the secondary binding events, respectively, *P*_T_ and *D*_T_ are the total concentrations of protein and DNA, respectively, *K*_1_ and *K*_2_ are the equilibrium dissociation constants for the primary and the secondary binding events, respectively, and *L* is the homodimerisation equilibrium constant. The equation was derived as shown in the Supplementary Material. All data fitting was performed using nonlinear least squares regression analysis implemented in Prism 9 (GraphPad).

## RESULTS

### Selection of residues for mutation

Previous research, including the published X-ray crystal structure for the CREB-CRE complex, has explored a _285_bZIP peptide corresponding to the fragment of CREB starting with K285 and continuing to K339 (two residues before the natural C-terminus of the protein) (55). Such a fragment contains the whole of the leucine zipper (LZ) motif, as well as the basic region (BR) and is capable of tight binding to the cognate CRE site characterised by low-nM *K*_D_ values. The X-ray structure analysis (PDB ID: 1DH3) confirmed that the most N-terminal residue directly involved in protein-DNA interactions is R286, and that the helical structure of DNA-bound CREB extends over the 285-334 range (46). In performing a Φ-value analysis, it is crucial that the mutations studied do not perturb the binding mechanism (10, 62). At the same time, examining as many positions as possible is needed to overcome the uncertainties in interpretation of individual Φ values and substantiate the results. A careful inspection of the X-ray structure of the _285_bZIP-DNA complex revealed that the side chains of the majority of residues that lie within the BR (K285-C310) are involved in the interactions with either DNA or Mg^2+^. The only residues with solvent-exposed side chains are K285, E287, V288, M291, E295, E299, R302 E306 and K309 (Figure 1A, B). K285 was excluded from analysis given its proximity to the N-terminus. The remaining eight residues were subjected to alanine-glycine (Ala-Gly) scanning, a well-established strategy to probe helix formation that minimises solvation effects of substituting polar amino acids (10–12, 14–16, 20, 21, 63, 64). Each of the solvent-exposed residues was mutated to both Ala and Gly, and the alanine mutant served as a pseudo-wild type to which the corresponding glycine mutant was compared.

### The basic region Is disordered but displays limited residual helical propensity

The ensemble of the unbound state in coupled folding and binding processes can contain elements of residual secondary and tertiary structures. The helical predictor AGADIR (65) suggests low levels of helical structure may exist within the DNA binding region of CREB, particularly residues 294-306 (Supplementary Fig 2, black lines). Circular dichroism (CD) measurements estimate the fractional helicity (FH) of the wild-type _285_bZIP is around 62%, whereas the LZ only constitutes 53% % of the residues (Supplementary Figure S3). Although CD is very good for observing trends it is notoriously difficult to estimate proportions of different secondary structure accurately. NMR studies can describe residual structure within the unbound state in a more accurate and residue-specific manner. We expressed and purified ^15^N and ^13^C/^15^N labelled versions of _285_bZIP and a related8omodimerizan-incompetent version with the leucine zipper removed, _285_BR. {^1^H}-^15^N HSQC spectra for the monomeric _285_BR showed narrow dispersion in the ^1^H^N^ dimension typical of a disordered protein whilst that for dimeric _285_bZIP showed more dispersion indicating some folded structure (Supplementary Figure S4). Here we calculate Secondary Structure Propensity (SSP) values for each residue based upon the method developed by the Forman-Kay laboratory (61). SSP values of 0 indicate a random coil structure, whilst values of 1 indicate helical structure. Monomeric _285_BR appears largely disordered but between 292-300 and 313-315 there are consecutive residues with SSP above 0.2 indicating low levels of helicity within those regions (Figure 1C, bright pink). We further investigated the mobility of the backbone in {^1^H}-^15^N heteronuclear nuclear Overhauser effect (NOE) experiments, and found the peptide to be highly dynamic, particularly towards the termini (Figure 1D, bright pink). The leucine zipper of _285_bZIP appears helical as anticipated, with some helix-fraying towards the C-terminus (Figure 1C, solid black). Interestingly, the SSP within the basic region is also increased considerably over the values found in the monomeric version. SSP in the basic region decreases almost monotonically with distance from the leucine-zipper. {^1^H}-^15^N heteronuclear NOE values decrease concomitantly with SSP in the same graded fashion with distance from the zipper indicating increased backbone dynamics – the rigid nature of the zipper itself is demonstrated by values above 0.65 for most of the leucine zipper, with increasing mobility towards the C-terminus where the helicity is lower. A four glycine residue (GGGG) insertion at position 307 (between the basic region and leucine zipper) within _285_bZIP completely abolishes this helical increase in the CREB dimer (Supplementary Figure S4). As a test of our SSP interpretations we also performed similar experiments for the _285_bZIP incubated with CRE DNA. Upon addition of CRE-DNA the SSP and {^1^H}-^15^N heteronuclear NOE values in the basic region increase still further, reflecting a fully folded and rigid helix which is consistent with the X-ray crystal structure for the CRE-bound complex (Figure 1C and 1D, dotted black). SSP and NOE results present a very consistent picture for all of the samples, indicating that as helicity is increased fast-timescale backbone dynamics are reduced, and vice versa.

**Figure 2.**
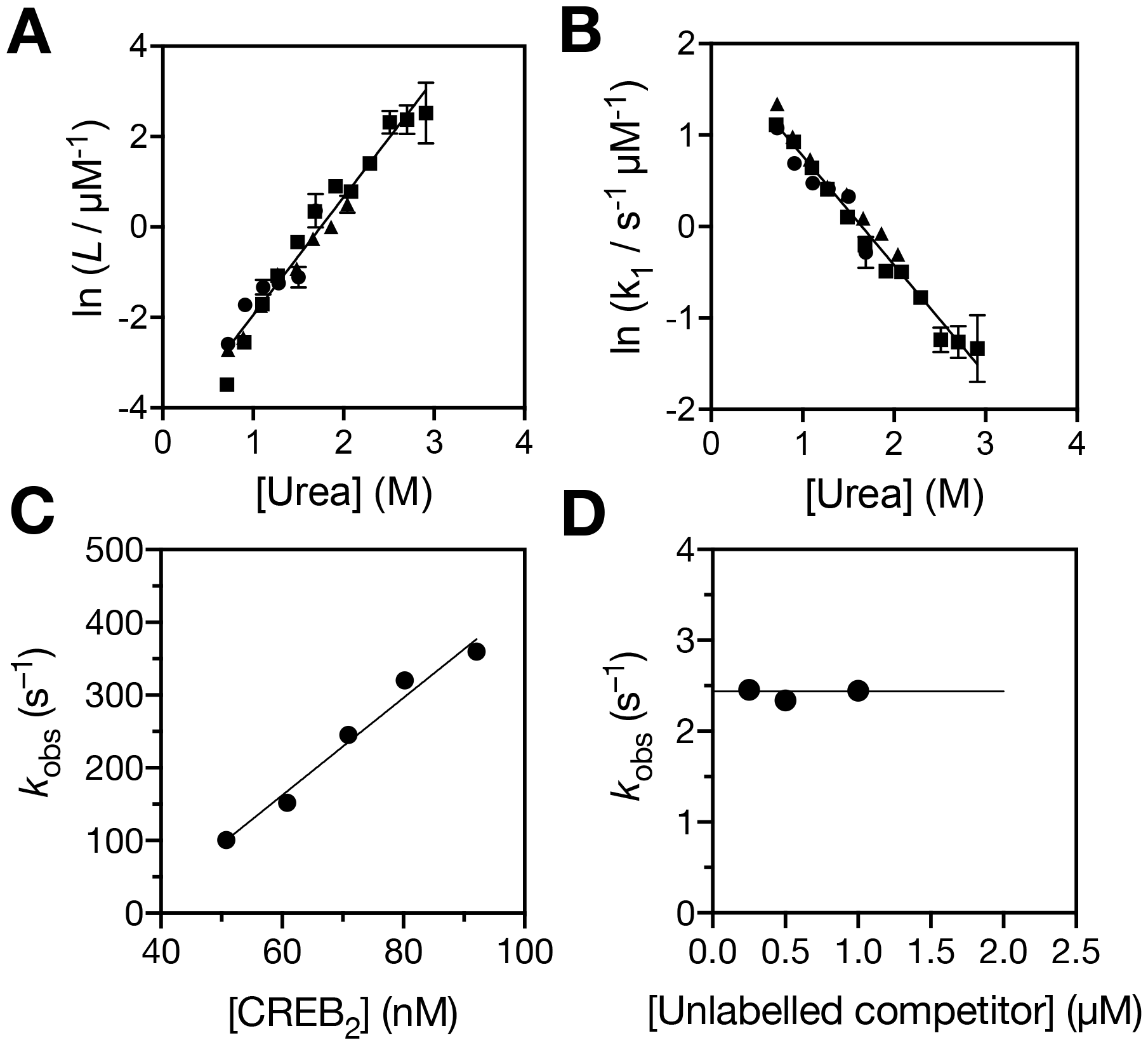
Characterisation of CREB binding interactions. Linear dependence of homodimerisation equilibrium constant (A) and dimerisation rate constant (B) on urea concentration. Extracted values from urea refolding stopped-flow kinetics studies with 2.2 uM (circles) 4.5 uM (triangles) and 9.1 uM (squares) CREB are in good agreement. Straight line fits enable extrapolation to estimate values in the absence of urea. C. Apparent association rate constant for CREB _285_bZIP with 10 nM AlexaFluor®488-CREh depends linearly on CREB _285_bZIP concentration. Concentration is for CREB dimer. Gradient of linear fit (solid line) represents *k*_on_. C. Apparent dissociation rate constant of CREB _285_bZIP from AlexaFluor®488-CREh is independent of concentration of out-competing unlabeled CRE DNA. Average of values represents *k*_off_.

### CREB bZIP forms tight homodimer complexes

We characterized the homodimerisation process using urea refolding kinetic studies to clarify the stoichiometry of CREB in the starting mixtures of CRE binding studies. We were fortunate to be able to make use of endogenous tyrosine fluorescence as a probe of homodimerisation. CREB was incubated in 8 M urea to favour monomer, and then homodimerisation was promoted by rapid dilution to reduce urea concentrations. Kinetic traces were well fit by an equation (Equation 1) derived for a simple 2-state reversible process 2*P* ⇋ *P*_2_ (Supplementary Figure S5). In further support of the model the extracted homodimer equilibrium dissociation constant (*L*) and folding rate constant (*k*_1_) both displayed a linear dependence upon final urea concentration that were independent of the CREB concentration (Figure 2A and 2B). Straight line fits of these data allow estimation of the binding affinity (11 ± 2 nM) and folding rate constant (7.1 ± 0.5 mM^-1^s^-1^) in the absence of urea. The gradients of the lines, or m-values (1.55 ± 0.07 and 0.71 ± 0.03 kcal mol^-1^ M^-1^), reflect changes in the solvent accessible surface area between the monomer and the transition state/dimer (66). The ratio of these two values (0.46) suggests that the transition state is roughly halfway in terms of the folding process.

### Wild-type CREB bZIP associates rapidly with CRE DNA

The binding of _285_bZIP to its cognate site was assessed using a fluorescently-labelled self-annealing oligonucleotide containing a single CRE motif from somatostatin promotor (55, 67). Upon rapid mixing of wild-type _285_bZIP with the AlexaFluor®488-CREh DNA by stopped-flow spectroscopy the fluorescence increased. We utilised total CREB concentrations above 100 nM in CREh association studies to probe interactions of dimeric CREB with CRE. Association kinetic traces collected under pseudo-first order conditions fit well to a single exponential decay function (Supplementary Figure 6A), with the observed rate constant being linearly dependent on protein concentration (Figure 2B). These observations are consistent with a simple 2-state process for binding of CREB and CREh, without the requirement for dimerisation, which is consistent with the majority of CREB being dimeric prior to mixing. The gradient of a straight line fit to the data, (6.7 ± 0.6) × 10^9^ M^−1^ s^−1^, was identified as the fundamental association rate constant (*k*_on_) for dimeric _285_bZIP with its target CREh.

To estimate the dissociation rate constant (*k*_off_) we performed a series of out-competition experiments where pre-formed AlexaFluor®488-CREh. _285_bZIP complexes were rapidly mixed with excess unlabeled CRE DNA. All traces fit well to a single exponential decay function (Supplementary Figure 6B), and the observed rate constants were independent of competitor concentration (Figure 2C), which allowed averaging to estimate *k*_off_ as (2.44 ± 0.03) s^−1^. When combined with our *k*_on_ estimate this indicates very tight binding of CREB dimer with CRE (*K*_D_ = (0.36 ± 0.04) nM).

### Mutations in the basic region of _285_bZIP have a marginal impact on residual structure

All substitutions introduced to the BR of _285_bZIP were predicted to have very marginal effects on helicity by AGADIR (Supplementary Figure S2). However, the sequence-based predictions do not take into account folding within the BR upon dimerisation that was observed by NMR. We used CD measurements to estimate the effect of mutation on helicity (Figure 3, Supplementary Figure S3). The FH of the wild-type _285_bZIP is 62%. The alanine substitutions either had no effect or led to a minor decrease in helicity, down to 56% in the case of R302A. Expectedly, all glycine mutants were less structured than the corresponding alanine mutants. The K309G mutant showed a 14% reduction in FH compared to K309A; for the rest of the substitutions the reduction was under 10%. The difference in FH between alanine and glycine mutants exhibited a clear upward trend in the direction of the C-terminus of the BR.

**Figure 3.**
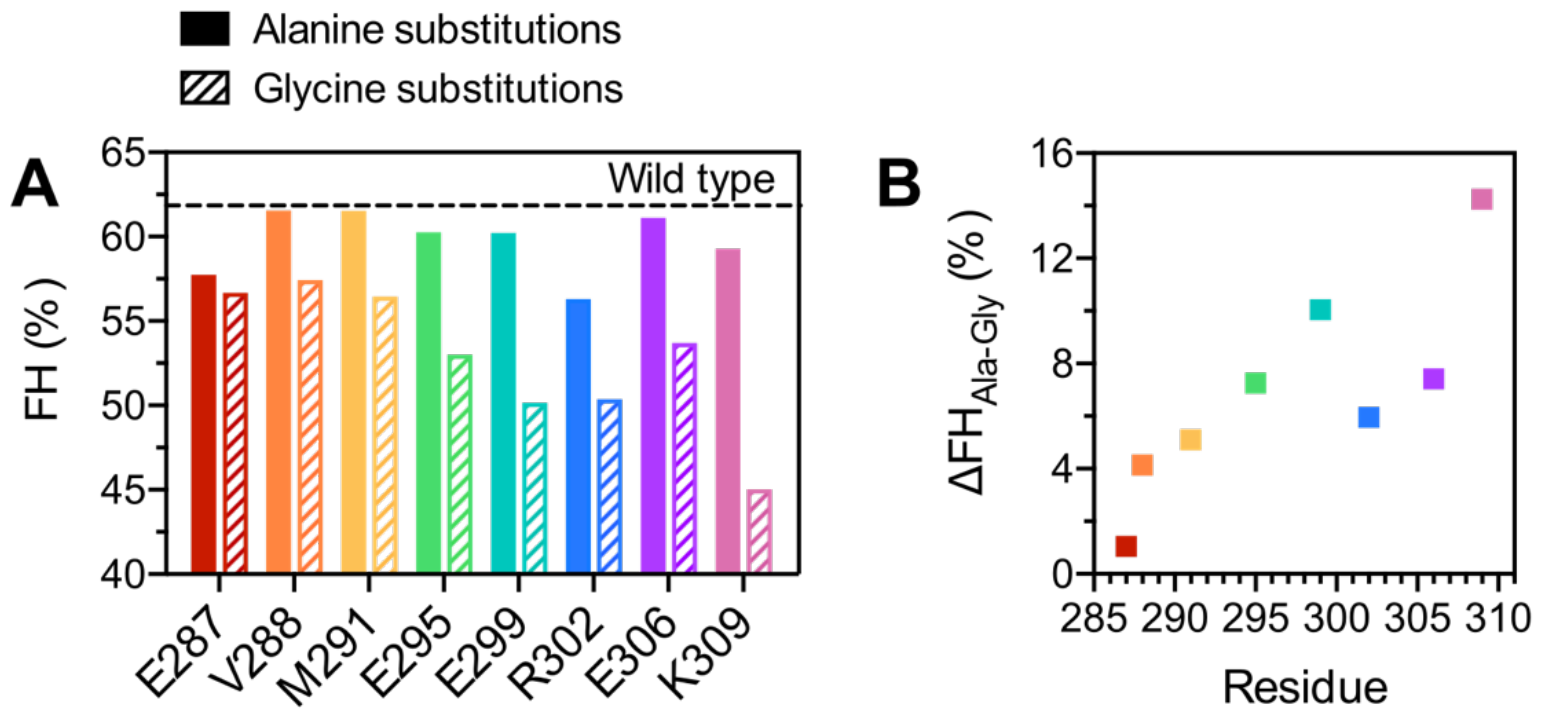
Effects of mutations on residual helicity of CREB _285_bZIP. A. Fractional helicities (FH) of the wild-type CREB _285_bZIP and its mutant versions estimated from the CD response at 222 nm. All alanine mutants are more structured than the corresponding glycine mutants. However, for all the residues except K309, the difference is within 10% of FH. B. The difference in FH between alanine and glycine mutants increases towards the C-terminus of the BR.

### Mutation modulates *k*_off_ rather than *k*_on_

Dissociation rate constants for the mutant versions of _285_bZIP were estimated as described for _285_bZIP, using a 40-fold excess of unlabelled competitor. Most of the alanine substitutions had a considerable effect on dissociation rates (Figure 4A, Supplementary Table 1). Mutations of residues at either ends of the BR resulted in stabilisation of the protein-DNA complex as opposed to the destabilising mutations in the middle of the BR. The only deviation from this pattern was the R302A substitution that demonstrated a remarkable 25-fold increase in *k*_off_ compared to the wild type. All glycine mutants had higher *k*_off_ values than the corresponding alanine mutants (Figure 4A, Supplementary Table 1). The ratio *k*_off,gly_ /*k*_off,ala_ correlated well with the difference in FH except for residue K309 (Figure 4D).

**Figure 4.**
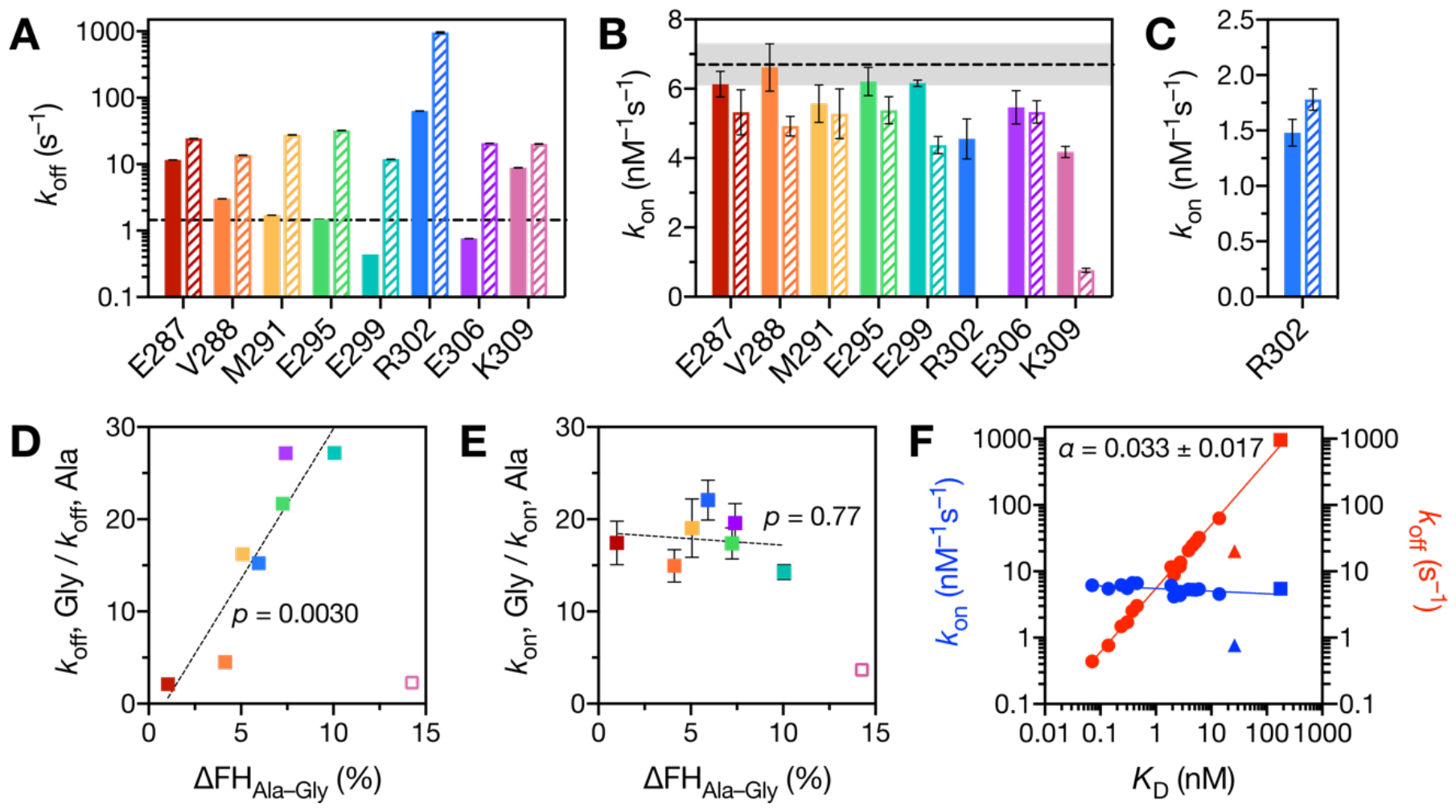
Kinetic effects of mutations in the BR of CREB _285_bZIP. A. Dissociation rate constants (*k*_off_). Alanine mutations are shown as solid bars, glycine mutations are shown as hatched bars, the wild type is shown as a dashed line. B. Association rate constants (*k*_on_) measured in stopped-flow experiments. Designations are identical to Fig. 4A. C. Association rate constants for R302A (solid bar) and R302G (hatched bar) mutants calculated as a ratio of *k*_off_ to the equilibrium binding constant (*K*_D_). The ratios of dissociation rate constants (D) and association rate constants (E) between each pair of glycine and alanine mutants plotted against the corresponding differences in FH. The decrease in residual structure upon glycine substitutions correlates with a decrease in stability of protein-DNA complex for every mutated residue except K309 (empty square). F. Linear free-energy plot showing *k*_on_ (blue) and *k*_off_ (red) rates against kinetic *K*_D_. The kinetic *K*_D_ for R302G (squares) was estimated as the equilibrium *K*_D_ multiplied by the ratio between kinetic and equilibrium *K*_D_ values for R302A. Outlier mutant K309G is shown as triangles.

Association kinetics of _285_bZIP mutants were analysed in the same manner as for the wild-type protein (Supplementary Figure 7) with a single exception. We were not able to directly measure the *k*_on_ value for R302G because of extremely low stability of its protein-DNA complex. Instead, we determined the dissociation constant (*K*_D_) by equilibrium titration (Supplementary Figure 8, Supplementary Table 1) and calculated *k*_on_ as a ratio of *k*_off_ to *K*_D_. Should the reaction involve a populated intermediate such an estimate may not match the *k*_on_ value measured in a stopped-flow experiment, although it can be very close depending on the individual microscopic rates. To mitigate this possibility, we employed the same method to estimate *k*_on_ for the R302A mutant and used it for comparison with the *k*_on_ of R302G.

The association rate constant of _285_bZIP with its target was largely unaffected by alanine mutations (Figure 4B, Supplementary Table 1). The substitutions of R302 and K309 led to a modest decrease in *k*_on_ (ca. 1.5-fold), however, the *k*_on_ value did not increase upon the substitutions of negatively charged residues E287, E295, E299 and E306. The glycine mutants appeared to exhibit either the same or marginally slower association rates than the corresponding alanine mutants, with a prominent exception of K309G that resulted in a 5-fold decrease in *k*_on_ compared to K309A (Figure 4B, Supplementary Table 1). Unlike with *k*_off_ the ratio *k*_on,gly_ /*k*_on,ala_ did not correlate strongly with the difference in FH (Figure 4E).

### The transition state is characterised by low Φ values

It is informative to consider changes in rate constants in the context of the overall change in stability. Fluorescence-based equilibrium titrations of _285_bZIP with CREh did not yield reliable *K*_D_ values due to the extremely high affinity. Direct measurements of *K*_D_ were possible for R302A and R302G, which both exerted a strong destabilizing effect as described above (Supplementary Figure 8). For the other mutants we used a kinetic estimate determined from the ratio of *k*_off_ to *k*_on_. As can be seen from a linear free energy relationship (LFER) plot, the effect of mutations on the stability of protein-DNA complex is almost entirely governed by dissociation rate (Figure 4F). The Leffler *α*, calculated as a gradient of log(*k*_on_) vs. log(*K*_D_) (11), has an extreme near-zero value of 0.033 ± 0.017, indicating that the transition state far more closely resembles the unbound state than the bound state. The K309G mutant is a clear outlier in the analysis. Structure formation in the transition state was further examined in a more positional manner by calculating Φ-values (Equation 2). Eight Φ-values were determined in total; each glycine mutant was compared directly with the equivalent alanine mutant to specifically probe helix formation. All the Φ-values were low (-0.1 to 0.2), with an exception of K309 which had a value of 0.7 ± 0.4 (Figure 5A, B).

**Figure 5.**
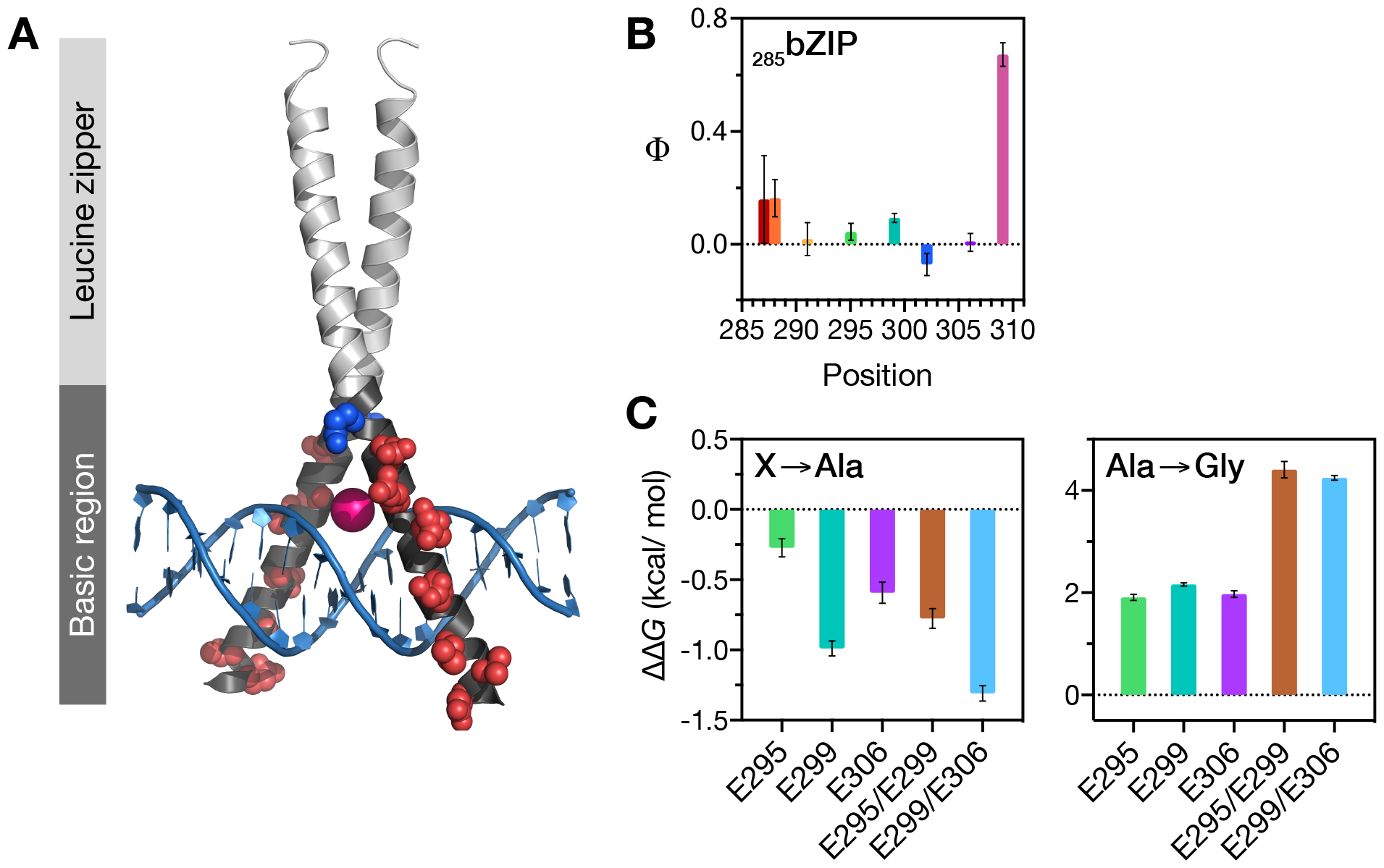
Φ-value analysis of the BR of CREB bZIP. A. Structural positions of residues with the determined Φ values (red spheres) in CREB _285_bZIP-DNA complex. The Mg^2+^ ion is shown as magenta sphere. Derived from 1DH3. B. Φ values calculated for alanine-to-glycine mutations in _285_bZIP. The near-zero numbers obtained for all analysed positions give evidence for a disordered transition state. C. The changes in binding free energy (ΔΔ*G*) upon X → Ala and Ala → Gly substitutions. The binding free energies to CRE were calculated from the *k*_off_ and *k*_on_ values. The effects of Ala → Gly substitutions on binding to CRE DNA are additive.

### Double mutations have additive effects on the stability of protein-CRE complex

We selected three residues (E295, E299 and E306) whose Ala-to-Gly substitutions produced the highest changes in binding free energy and analysed the double mutant versions of _285_bZIP with either E295/E299 or E299/E306 being swapped for alanines and glycines. The double glycine mutants appeared to exhibit roughly the same *k*_on_ rates as their double alanine counterparts (Supplementary Figure S9A-B, Supplementary Table 1). However, for both E295G/E299G and E299G/E306G, we observed a more than three orders of magnitude increase in *k*_off_ compared to E295A/E299A and E299A/E306A, respectively (Supplementary Figure 9C, Supplementary Table 1). As a result, the ΔΔ*G* values for double Ala-to-Gly substitutions determined from *K*_d_ comparisons were exceeding 4 kcal/mol, which is approximately the sum of the ΔΔ*G* values determined for the corresponding single mutations (about 2 kcal/mol each) (Figure 5C). Both E295/E299 and E299/E306 yielded near-zero Φ values (0.03 ± 0.04 and −0.022 ± 0.012, respectively).

## DISCUSSION

### Wild-type CREB kinetic and equilibrium parameters

The bZIP domains are observed to fold upon binding their cognate DNA sequences (2, 43, 68–70). We have examined the interaction of the isolated bZIP domain of CREB with CRE and find the binding affinity is sub-nanomolar, which is slightly lower than previous estimates for the full-length protein of around 1–2 nM (52, 71), but in line with a recent 1 nM upper limit (72). Our results rationalise the observed differences in terms of the experimental approach. The equilibrium dissociation constant for homodimerisation, *L*, is around 10 nM and thus over 10-fold higher than the *K*_d_ for CRE binding. This means that equilibrium approaches to estimate *K*_d_, such as those in the older assessments, necessarily provide an *apparent* binding constant that involves a very significant contribution from dimerisation. In contrast our kinetic CRE binding experiments are performed at concentrations above *L* and so we probe the interactions of CREB dimer.

The nanomolar value of *L* is also noteworthy because previously reported values differ wildly from tens of µM (73, 74) to sub-nanomolar (75). Our value is similar to that found by Bentley et al. for full-length CREB (72) who suggest CREB may be largely dimeric within the cell. For reference cellular CREB concentrations are around 100-fold higher than our reported homodimerisation *K*_D_ (76, 77). In determining *L* we revealed that homodimerisation was well described as a 2-state process without populated intermediates. The folding rate constant we obtained, of 7 x 10^6^ M^-1^s^-1^, is similar to that reported for GCN4 (17, 78), and fairly typical of diffusion-controlled protein-protein interactions for both folded and disordered proteins (79).

The kinetic rate constant for binding to CRE however, is much higher. The value obtained for *k*_on_, of around 7 x 10^9^ M^-1^s^-1^, is similar to those obtained *in vitro* for other transcription factors with their targets, including GCN4 bZIP (80). Such a high value, around that of the diffusion-limit for uncharged spheres (despite presumed activation energy barriers and geometric constraints), indicates substantial electrostatic rate enhancement. This is presumably a result of complementary charges for the bZIP (+8) and DNA (-30 for AlexaFluor®-488 CREh) providing long-range attractive forces. Apparently consistent with this, removal of a single positive charge by alanine mutation (R302A) does lower *k*_on_. Our observation that removing a single negative charge using the same strategy does not increase *k*_on_ correspondingly implies enhancement is not simply a reflection of the net charges of the molecules however. One of the most distinctive features of CREB bZIP is a conserved continuous stretch of five positively charged amino acids R301-K305 within the BR (Figure 1A). It is feasible this stretch may play a key role in the initial charge-driven association of protein and DNA via electrostatic steering. Other systems with very significant electrostatic rate enhancements, such as barnase-barstar, also contain highly complementary charged “docking” surfaces as well as high net charges (81).

Residency times for transcription factors on their target promoters must be within useful timescales for successful transcription within eukaryotic cells. Notably the value we obtain for *k*_off_, of around 1 s^-1^, is similar to the 0.42 s^-1^ at which individual fluorescently labelled CREB molecules were observed to leave their long-residency locations in mouse cortical neurons (82). These findings suggest that isolated bZIP domains can encapsulate much of the behaviour of the full-length protein.

### Helical structure is propagated into the basic region in the dimer

NMR analysis showed that higher helical propensities in the C-terminal end of the BR are not present without the folded LZ, and, within the dimer, SSP scores decrease almost linearly with distance from the zipper. Thus, the helical LZ stabilises helical structure in the neighbouring BR, with the stabilising effect diminishing with distance i.e. helicity is propagated outwards by dimerisation. Single-molecule FRET studies of CREB with fluorescent labels at either end of the LZ (residues 310 and 337) and BR (residues 270 and 310) are consistent with dimerisation leading to structural change in the ensemble of the BR (72). Changes to helicity observed by CD for mutant proteins can add further detail. The decrease in FH between each alanine mutant and its corresponding glycine mutant is almost linear with the residue position (Figure 3B) in striking agreement with the spatial pattern of helicity of the apo-state using the more robust NMR method. However, mutation to alanine, a helix favouring residue, causes an appreciable reduction in FH for two residues; E287 and R302. E287 is in a good position to stabilise the helix since its negative charge could offset some of the positive charge towards the N-terminal end of the helix (which would generally interact negatively with the helix dipole). Helix N-capping residues have been demonstrated in GCN4, and suggested within roughly half of bZIP basic regions (83). The crystal structure of the CRE-bound complex shows R302 forming putative salt bridges with E299 and E306 (46), which if present in the unbound state would undoubtedly stabilise helical structure.

Our NMR and Ala-Gly scanning results challenge the concept of a boundary between the BR and LZ in CREB. Historically a linker region between BR and LZ has sometimes been defined in bZIPs (84, 85), and helical predictions by AGADIR often show a dip in helicity between the DNA binding region and zipper domain as we observe in the prediction for CREB (Supplementary Figure S2). However, NMR and Ala-Gly scanning have uncovered here an allosteric-like propagation of structure from the adjacent dimerisation domain in the dimeric CREB bZIP. This finding implies that the sequence of bZIP “linkers” could be functionally very important even when it makes no intermolecular contacts since we reveal that binding affinity depends upon pre-existing helicity propagated through the linker.

### The transition state strongly resembles the unbound state

To gain additional insight into the folding and binding mechanism we used alanine and glycine mutants of CREB bZIP in comparative kinetic analyses. Despite binding affinity varying by more than 3 orders of magnitude between mutants, *k*_on_ values vary very little, with all but one mutant being in the range of 4–7 x 10^9^ M^-1^s^-1^. Instead, there is a very clear and substantive (up to 30-fold) increase in *k*_off_ upon glycine mutation for each Ala-Gly pair (Figure 4A). With the notable exception of K309, there is a good linear correlation between the relative *k*_off_ and change in FH. Thus, stabilising helical conformations in the BR leads to tighter binding largely through changes in *k*_off_. Indeed, the low Φ values across almost the entire length of the BR (E287–E306) and a near-zero Leffler α imply that the transition state is structurally early and resembles the unbound protein (Figure 4F and Figure 5). The K309G mutant is an outlier in that it does not lie on the straight line revealed on the LFER plot, and the resulting Φ-value is considerably higher than the others i.e. this residue is in a largely helical conformation at the transition state. Interestingly our NMR investigation of CREB bZIP indicates that (in the absence of mutation) K309, identified by Uniprot as the final residue of the basic region, is actually largely helical in the unbound state. Given the observed single higher Φ -value close to the LZ, and distance-dependent stabilisation of BR structure by the helical LZ it is tempting to speculate that folding may initiate from the LZ with further helix formation being stimulated towards the N-terminus in a downhill fashion. The overall level of residual helicity in the BR varies across the bZIP family (43), but this is plausibly a common mechanism for all bZIP dimers. Robustelli *et al*. have previously speculated that GCN4 folding closer to the LZ might stimulate folding further away (86), however this has not been tested.

Φ-value analysis is currently the only experimental method capable of probing the structure of folding transition states, yet it is sometimes criticised because making mutations can make it difficult to adhere to the basic assumptions of the underlying theory. Analysis assumes that mutations introduced to probe the transition state have a negligible effect on the unfolded state (62). To better comply with that requirement, it is ideal to study an IDP with as little residual structure as possible, so that substitutions minimally impact the disordered state of the protein. Our CREB construct does not have high levels of helical structure, but mutations made within the basic region in this study could, and do, alter the unbound (disordered) state somewhat. However multiple lines of evidence point against de-routing effects; changes in protein-DNA complex stability are additive, Φ-values remain consistent when calculated in the context of single and double mutants, and the LFER plot is linear (62). Alanine substitutions appeared to produce rather small changes in binding free energy compared to the wild-type _285_bZIP (< 1 kcal/mol in magnitude) which also complies with the Φ-value requirement for perturbations to be small. Taken together, the _285_bZIP peptide maintains all native protein-DNA interactions while being a convenient target for the Φ-value analysis and therefore represents a good model for studying the mechanism of DNA binding by CREB. Our results are robustly consistent with little to no folding (gain in helical structure) between the unbound and transition state ensembles for folding and binding of CREB with its DNA target CRE.

Most studies for the interaction of the IDPs with protein partners have observed a relatively unstructured transition state. They have proposed either induced fit or a mechanism combining the features of both induced fit and conformational selection (6, 7, 11–13, 18–20, 87–93). This study explores the mechanism of coupled folding and binding with a DNA partner instead, however our results echo the findings with protein partners in having a very early transition state that strongly resembles the unbound state. Since the transition state must be formed after the encounter complex this indicates a heterogenous and dynamic encounter complex as well. This could plausibly facilitate successful and efficient rearrangement from the encounter complex.

### Fishing for fly-casting

One stated advantage to protein disorder is that it may enhance association rate constants over those for folded proteins through a mechanism known as “fly-casting” where encounter rates are accelerated by increased protein radii (9). The effect has been implicated in molecular dynamics simulations (94–99), but despite its continued prominence in literature discussion direct experimental evidence for this fly-casting effect is lacking. Enhanced association rates have been observed for mutants of a major histocompatibility complex-like protein, MICA. MICA is mostly folded in its apo-state but has a disordered segment which folds into two helix turns and a loop upon binding its immunoreceptor NKG2D (100). Point mutations aiming to destabilise pre-existing structure of the disordered segment (by weakening stabilising MICA self-interactions) lead to tighter binding, with 3-fold enhanced *k*_on_ for charge-neutral mutations Q120I and N68W. This could reflect a fly-casting effect, however the mutations are very non-conservative and it may instead reflect destabilisation of the unbound state, or a decreased energetic cost for rearrangement from the encounter complex to the transition state. A more targeted experimental investigation is notable in its absence. A capillarity theory for fly-casting revealed that fly-casting is more likely when the protein unfolding barrier is small, the binding affinity high, and the chain can be somewhat rigidly extended towards the target i.e. they suggest a role of residual structure in the unbound state (97). The system investigated here, which meets all these criteria, is therefore an excellent candidate for observing fly-casting experimentally. Indeed the papers outlining the theory highlight protein-DNA binding, contain supplementing simulations of an operator binding to DNA (9, 97), and even mention bZIP domains by name (9). Whilst remaining dynamic the BR of CREB has high charge content, and even significant helical structure propagated outwards from the zipper. It has all the hallmarks of an effective fly-casting scenario.

A **Φ** -value like approach to revealing the fly-casting phenomenon (where stability is altered but the binding interface remains unchanged) has been suggested several times (9, 97, 98). In this work we have done precisely this. Ala-Gly scanning enables us to consider specifically how changes in helicity affect binding kinetics. We have systematically decreased the helical propensity through glycine mutation but we have not observed an increase in *k*_on_ as the fly-casting model would predict. A single glycine mutant (out of eight), R302G, had a higher *k*_on_ than its alanine counterpart. However, *k*_on_ was calculated indirectly in this case due to the extreme destabilisation resulting from glycine mutation, and the apparent increase of (1.20 ± 0.12) -fold is relatively small. Furthermore, given the number of measurements, one to three values outside the standard error are to be expected statisticall. All other glycine mutants had marginally smaller or similar *k*_on_ values to their alanine counterparts. Thus, we have not observed an enhancement of binding rates by protein disorder experimentally.

In molecular dynamics simulations Huang et al. did observe higher association rates for more disordered versions of pKID (an activation domain of CREB) but this was due to a reduced energy barrier to the transition state (101). Recent atomistic MD simulations for another induced fit process also found more disordered encounter complexes were more likely to proceed to bound state (88). In simulations more ordered ligands in encounter complexes have been observed to unfold and refold again during the process of attaining the bound state (88), and there is some limited evidence that disordered proteins may typically have high k_on_ even in the absence of electrostatic rate enhancement (102). However any requirement for structure in the transition state, possibly even in induced fit processes, could likewise disadvantage disordered proteins (8). The balance between these two requirements will likely depend upon the individual macromolecules, the nature of the binding surface, and the required final topology, and eclipse any kinetic advantage from fly-casting. The predicted enhanced capture rates from fly-casting are only modest, perhaps 1.4–1.6 fold (9), so any effect would be easily overshadowed by energetic terms contained in an exponential. These terms could be reduced for disordered proteins if flexibility is important for rearrangements to reach the transition state from the encounter complex.

Unlike the simulations we are not able to make comparisons with a “fully folded” version. Our not observing a rate enhancement could potentially be because the changes made in “foldedness” are too small or localised. CD spectra indicate the loss of helical structure from glycine mutation is equivalent to up to around 20% of FH in the BR, however this is actually a considerable decrease considering it is only partially helical initially. Although we are not able to completely rule out the existence of a fly-casting effect our not observing one in this “model” system with the suggested experimental approach questions its practical relevance. Ultimately what Biology will care about is the rate constant(s), and this is determined by the concentration of the transition state rather than the encounter complex – transition state and electrostatic effects must be expected to dominate over any putative fly-casting in most circumstances.

## CONCLUSIONS

Application of Ala-Gly scanning to the folding of CREB upon CRE binding has revealed CREB to have a transition state strongly resembling the structural ensemble of the free protein. The results are consistent with an electrostatically-enhanced binding step followed by fast downhill folding propagating from the already helical leucine zipper.

## DATA AVAILABILITY

NMR assignments are detailed in BMRB depositions 50880, 50872, 50873.

## SUPPLEMENTARY DATA

Supplementary data include Supplementary Methods (equation derivations), nine supplementary figures, and one supplementary table.

