## Supplementary Materials for "The transition state for coupled folding and binding of a disordered DNA binding domain resembles the unbound state"

### SUPPLEMENTARY TABLES

| Construct | $k_{off}$ ($s^{-1}$) | $k_{on}$ ($s^{-1}nM^{-1}$) | $K_{D, CRE}$ (nM) | $K_{D, control}$ (nM) |
|---|---|---|---|---|
| Wild type $_{285}$bZIP | 2.531 ± 0.006 | 6.7 ± 0.6 | | 88 ± 6 |
| $_{285}$bZIP E287A | 11.59 ± 0.05 | 6.1 ± 0.4 | | 51 ± 5 |
| $_{285}$bZIP V288A | 3.022 ± 0.009 | 6.6 ± 0.7 | | 94 ± 7 |
| $_{285}$bZIP M291A | 1.708 ± 0.004 | 5.6 ± 0.5 | | 85 ± 6 |
| $_{285}$bZIP E295A | 1.481 ± 0.003 | 6.2 ± 0.4 | | 43 ± 4 |
| $_{285}$bZIP E299A | 0.4379 ± 0.0007 | 6.16 ± 0.09 | | 41 ± 4 |
| $_{285}$bZIP R302A | 63.2 ± 0.3 | 4.6 ± 0.6 | 41 ± 4 | 1200 ± 70 |
| $_{285}$bZIP E306A | 0.7596 ± 0.0012 | 5.5 ± 0.5 | | 44 ± 5 |
| $_{285}$bZIP K309A | 8.85 ± 0.04 | 4.18 ± 0.16 | | 320 ± 20 |
| $_{285}$bZIP E295A/E299A | 0.4874 ± 0.0008 | 4.8 ± 0.4 | | 21 ± 3 |
| $_{285}$bZIP E299A/E306A | 0.1411 ± 0.0002 | 3.40 ± 0.07 | | 26 ± 4 |
| $_{285}$bZIP E287G | 24.45 ± 0.13 | 5.3 ± 0.6 | | 45 ± 4 |
| $_{285}$bZIP V288G | 13.63 ± 0.08 | 4.9 ± 0.3 | | 123 ± 8 |
| $_{285}$bZIP M291G | 27.7 ± 0.2 | 5.3 ± 0.7 | | 184 ± 13 |
| $_{285}$bZIP E295G | 32.1 ± 0.2 | 5.4 ± 0.4 | | 112 ± 9 |
| $_{285}$bZIP E299G | 11.90 ± 0.05 | 4.4 ± 0.2 | | 130 ± 13 |
| $_{285}$bZIP R302G | 960 ± 20 | | 540 ± 30 | 3080 ± 150 |
| $_{285}$bZIP E306G | 20.64 ± 0.13 | 5.3 ± 0.3 | | 241 ± 13 |
| $_{285}$bZIP K309G | 20.18 ± 0.16 | 0.77 ± 0.06 | | 810 ± 50 |
| $_{285}$bZIP E295G/E299G | 649 ± 16 | 3.7 ± 1.0 | | 69 ± 7 |
| $_{285}$bZIP E299G/E306G | 214.1 ± 1.9 | 4.0 ± 0.3 | | 95 ± 9 |

Supplementary Table S1. Kinetic and thermodynamic parameters for the DNA binding of all examined $_{285}$bZIP and $_{277}$bZIP constructs. The errors represent the errors of the fit.

### SUPPLEMENTARY FIGURES

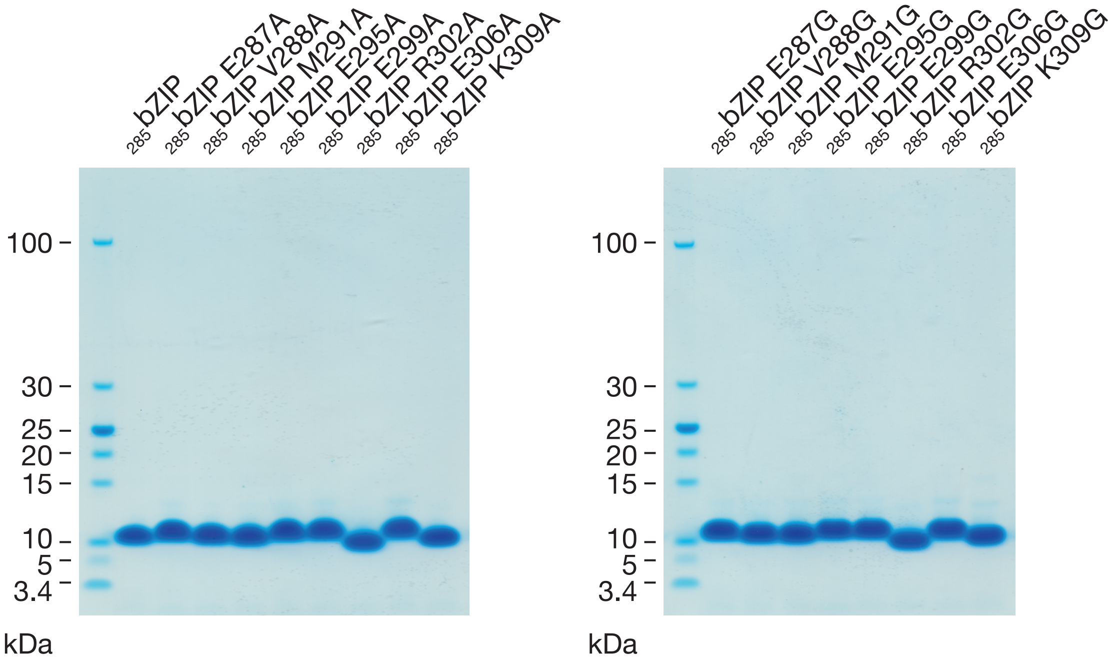


Supplementary Figure 1. SDS-PAGE analysis of CREB $_{285}$bZIP constructs. Molecular weights of the protein standards (first lane) are indicated. Expected molecular weight is 6.7 kDa. Note that disordered proteins typically bind less SDS than globular proteins as a result of their sequence compositions and therefore migrate more slowly in SDS-PAGE analysis[*]. Precise molecular weight was confirmed by mass spectrometry (ESI-MS). Mutation of positively charged residues R302 and K309 has a minor effect on migration.

---

[*] Vladimir N. Uversky and A. Keith Dunker, "Multiparametric Analysis of Intrinsically Disordered Proteins: Looking at Intrinsic Disorder through Compound Eyes", *Analytical Chemistry* **2012** *84* (5), 2096-2104

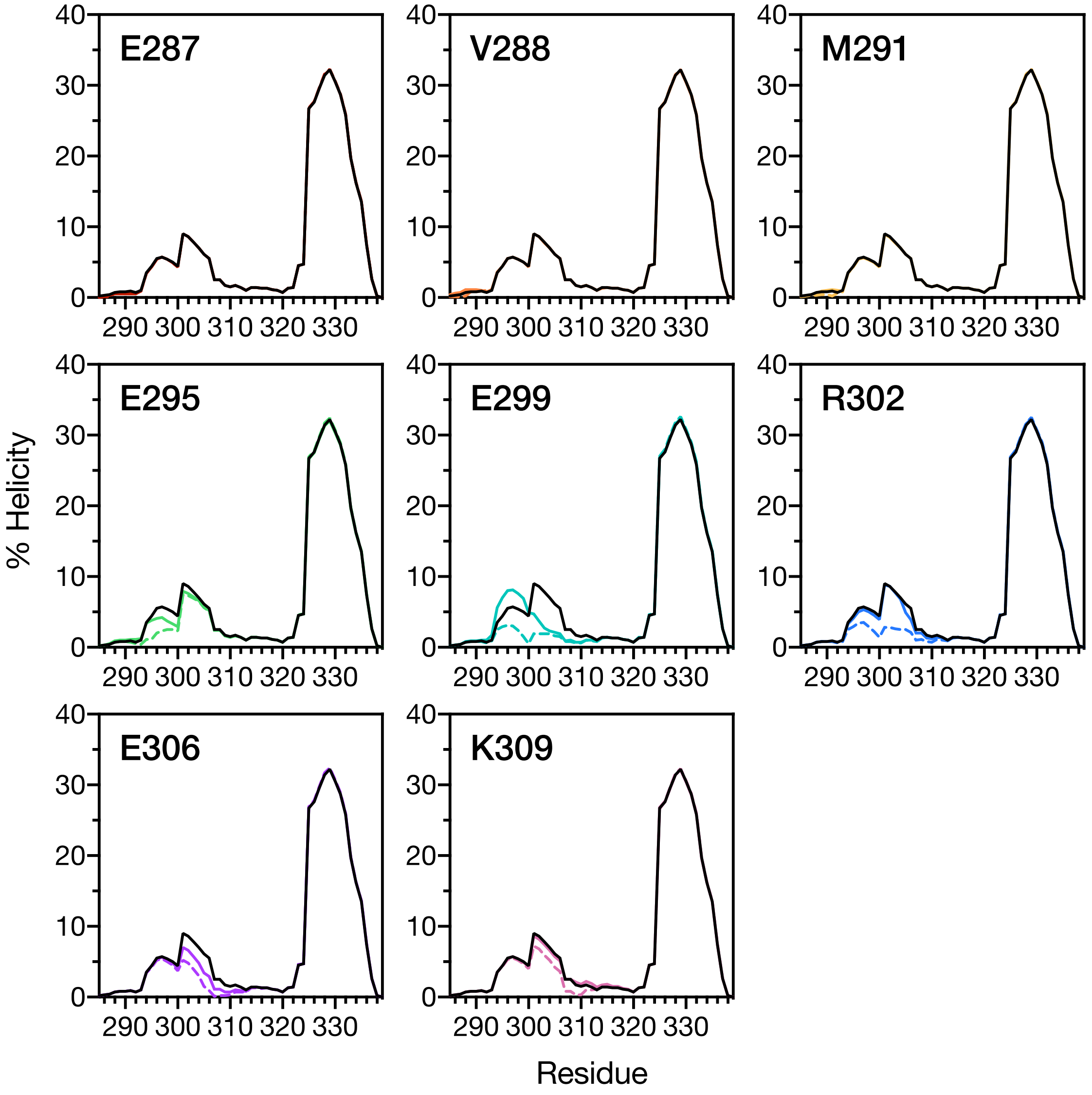


Supplementary Figure 2. AGADIR predictions of helicity based upon amino acid sequence. CREB $_{285}$bZIP sequence predictions are shown as black lines. Predictions for alanine mutants (solid lines) and glycine mutants (dashed lines) are shown in individual panels. Where coloured lines are not visible this is because predictions overlap with CREB $_{285}$bZIP.

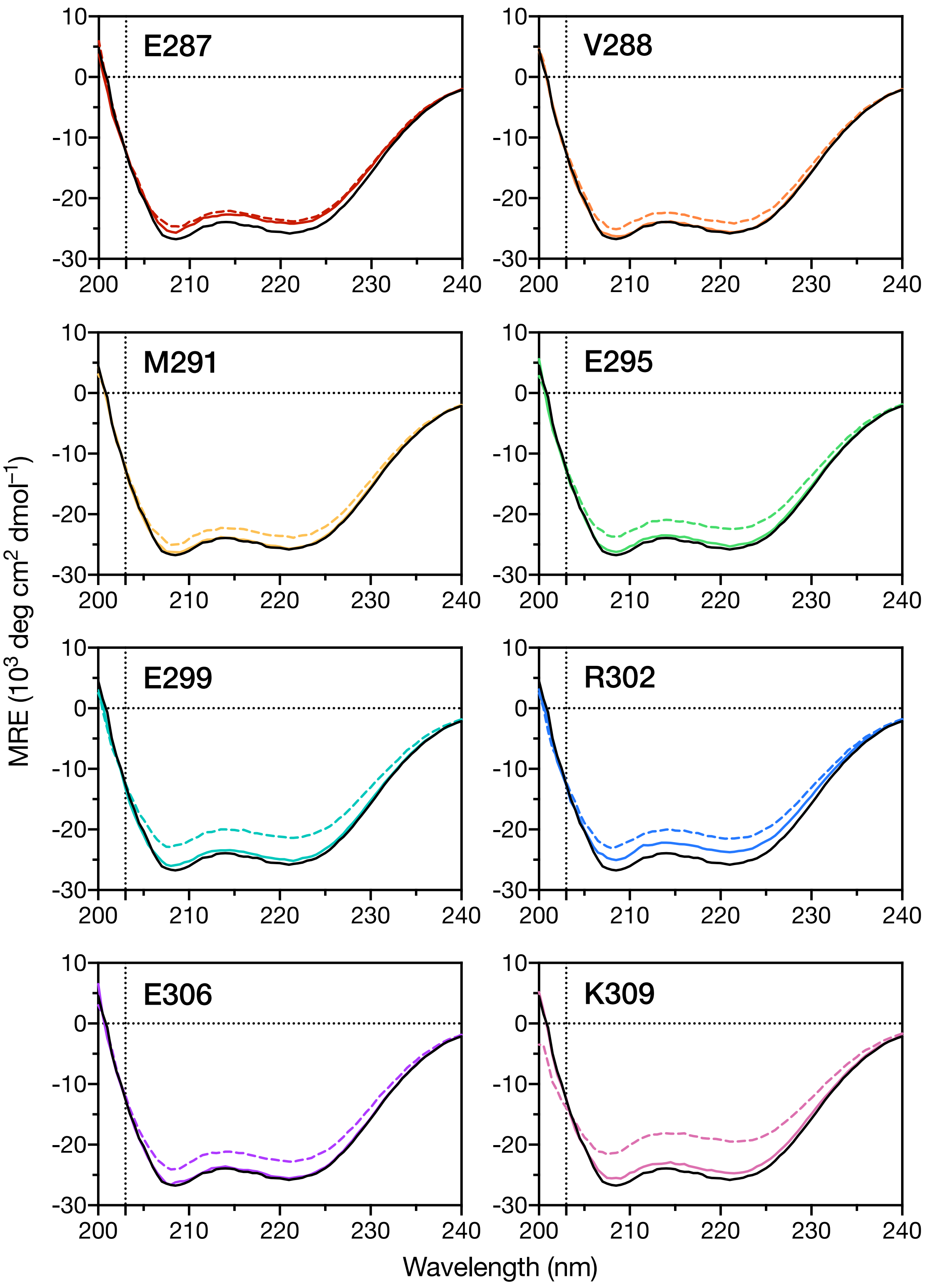


Supplementary Figure 3. Circular dichroism spectra for CREB $_{285}$bZIP and mutants. 20 µM protein samples in 10 mM MES pH 6.5, 150 mM NaCl, 10 mM $MgCl_2$, 0.05% Tween-20 (biophysical buffer) at 25 °C. CREB $_{285}$bZIP is shown as a solid black line, and alanine mutants (coloured solid lines) and glycine mutants (coloured dashed lines) are shown in individual panels. Spectra displayed are an average of three replicates. CD signal at 222 nm can be used as an indicator of helical structure.

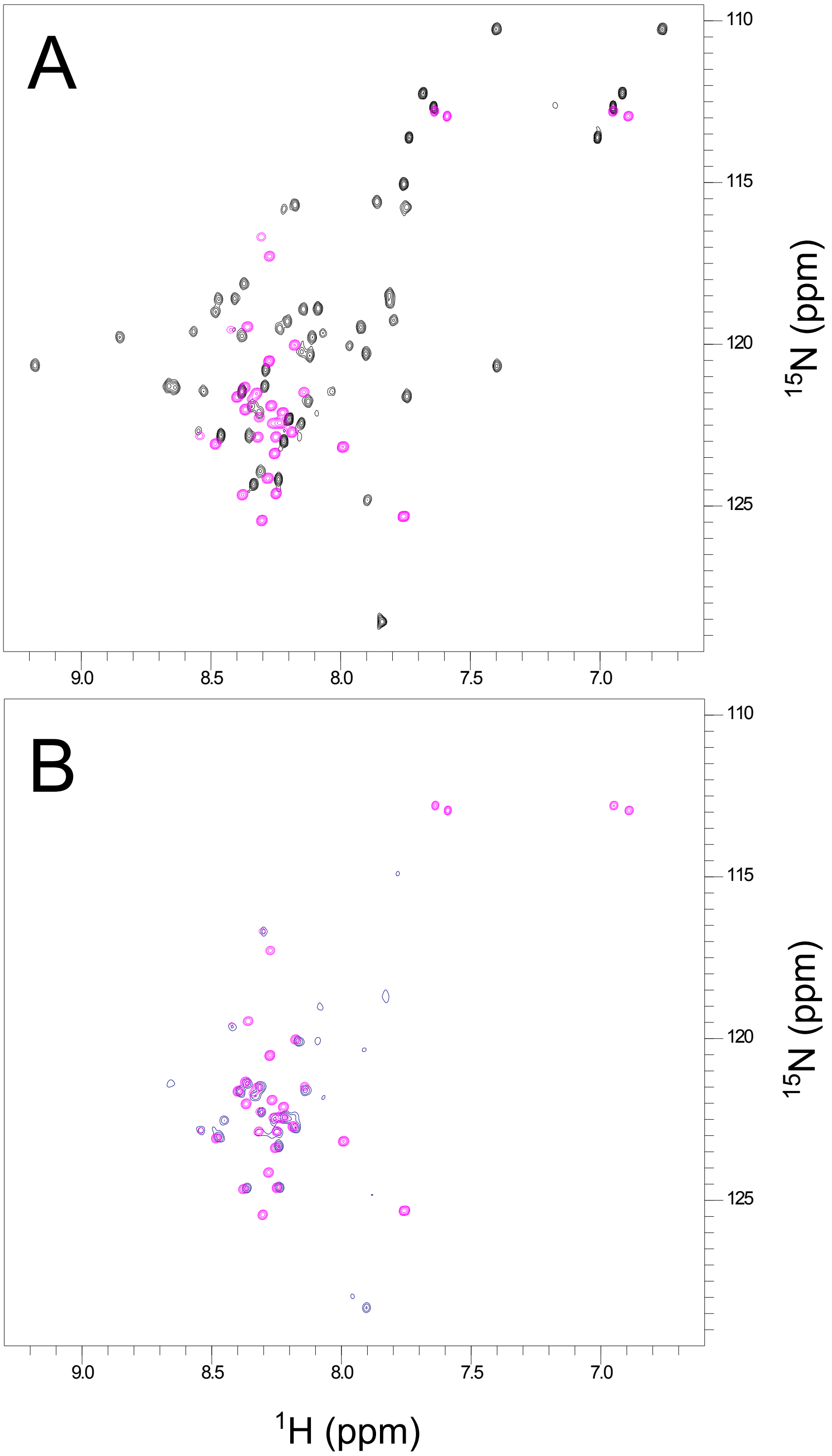


Supplementary Figure 4. Overlays of 600 MHz $^1H$-$^{15}N$ HSQC NMR spectra for CREB $_{285}$BR (magenta in A and B), CREB $_{285}$bZIP (black in A) and natural abundance HSQC spectrum of CREB $_{285}$bZIP-ins307_GGGG (navy blue in B) collected at 25 °C. Peak overlay indicates similar environments/secondary structure for the basic region residues of the GGGG insertion mutant with the monomeric basic region construct. Samples contained 0.3 -1 mM protein in 95% $H_2O$/5% $D_2O$ with 10 mM Tris-$d_{11}$ (Sigma), 150 mM NaCl, 10 mM $MgCl_2$, 1 mM $NaN_3$ plus protease inhibitors (Pierce$^{TM}$, Thermo Scientific).

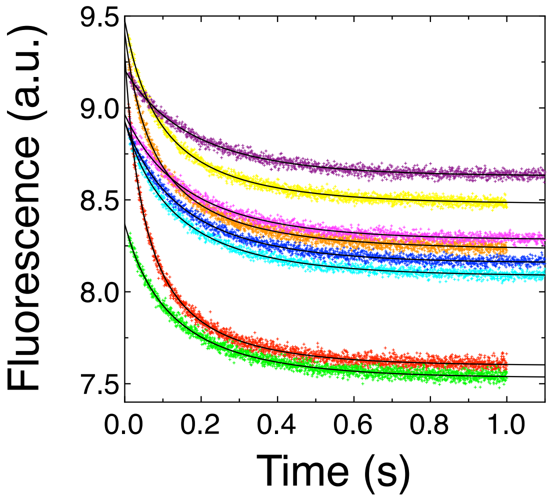


Supplementary Figure 5. Stopped-flow kinetic trace from urea refolding studies. CREB $_{285}$bZIP in 8 M urea was mixed rapidly in a 1:11 ratio to achieve final concentrations of 4.5 μM protein, 0.72 M urea (red), 0.89 M urea (orange), 1.08 M urea (yellow), 1.27 M urea (green), 1.48 M urea (cyan), 1.66 M urea (blue), 1.86 M urea (pink) and 2.04 M urea (purple). Buffers contained 10 mM MES pH 6.5, 150 mM NaCl, 10 mM $MgCl_2$, 0.05% Tween-20 (biophysical buffer) and measurements were performed at 25 °C. Black line is best fit to Equation 1.

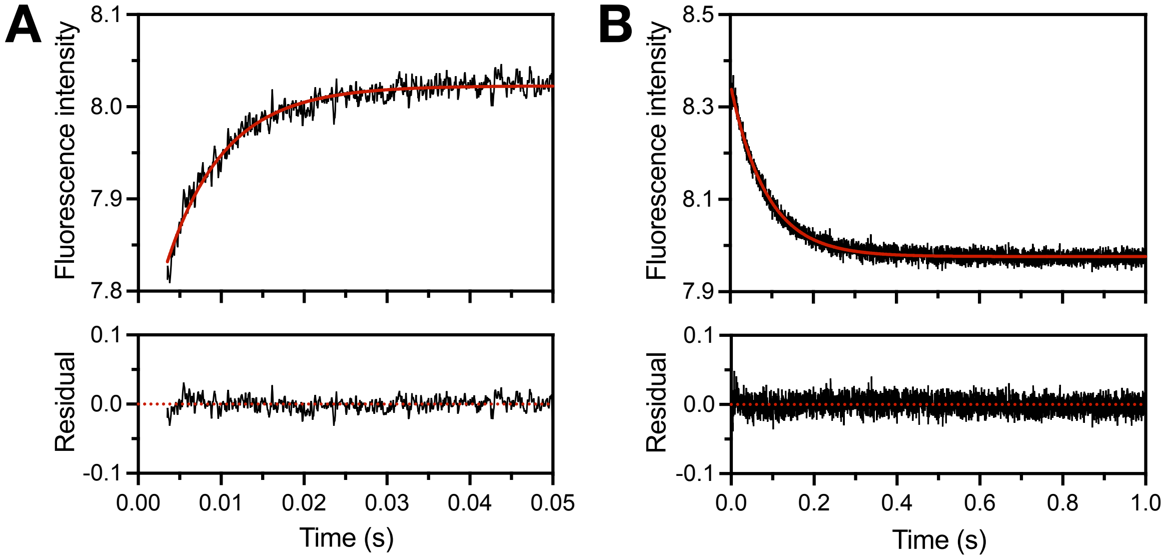


Supplementary Figure 6. Representative stopped-flow kinetic traces for CREB $_{285}$bZIP E287A association with (A) and dissociation from (B) AlexaFluor®488-labeled CREh. Kinetic data are shown as black lines and fitted single exponential decay functions as red lines. All solutions are in 10 mM MES pH 6.5, 150 mM NaCl, 10 mM $MgCl_2$, 0.05% Tween-20 (biophysical buffer) and measurements were performed at 25 °C. In (A) 10 nM AlexaFluor®488-labeled CREh is mixed rapidly with 200 nM CREB $_{285}$bZIP in a 1:1 ratio. In (B) a pre-equilibrated mixture of 10 nM AlexaFluor®488-labeled CREh and 100 nM CREB $_{285}$bZIP is mixed rapidly with 4 μM unlabelled competitor CRE DNA.

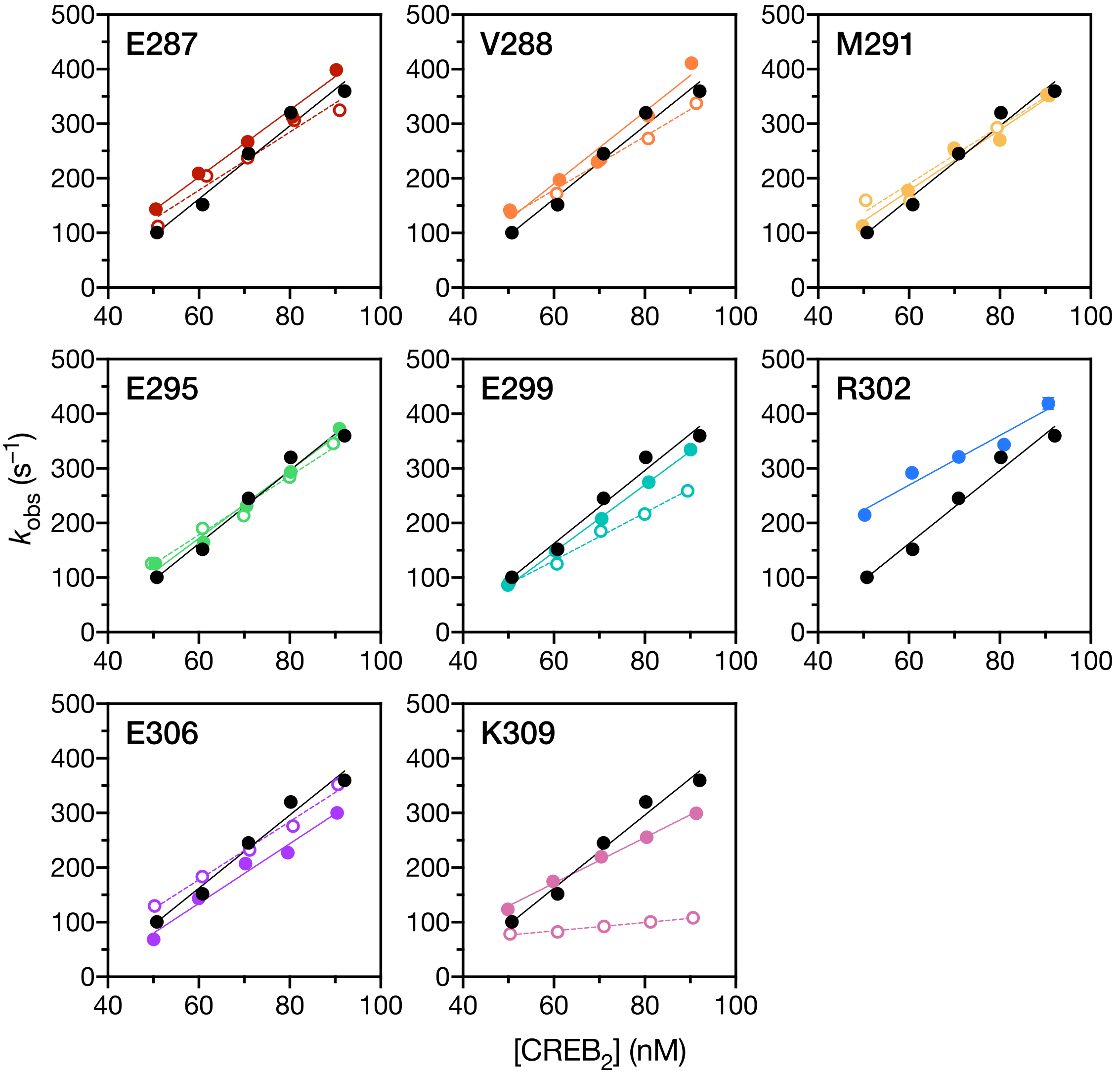


Supplementary Figure 7. Apparent association rate constants with 10 nM AlexaFluor®488-CREh. CREB $_{285}$bZIP is shown as a solid black circles, and alanine mutants (coloured solid circles) and glycine mutants (coloured open circles) are shown in individual panels. Concentrations are for dimeric CREB. Lines are straight line fits, and the gradient represents $k_{on}$. Kinetic stopped-flow data were collected in 10 mM MES pH 6.5, 150 mM NaCl, 10 mM $MgCl_2$, 0.05% Tween-20 (biophysical buffer) at 25 °C.

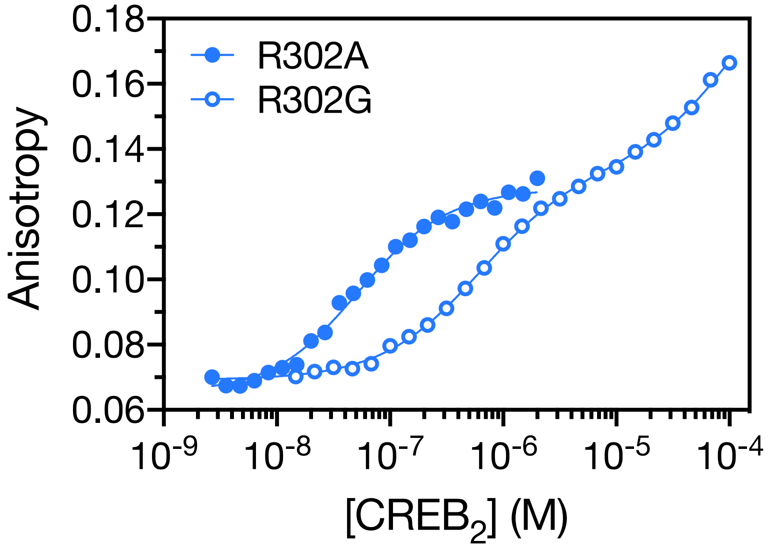


Supplementary Figure 8. Anisotropy equilibrium titrations for R302A (solid circles) and R302G (open circles) mutants. Protein concentration shown is monomeric concentration divided by two. At lower CREB concentrations monomeric CREB is not expected to be all dimeric so data are fit to Equation 2 which accounts for homodimerization and two binding events (solid lines). Concentration of AlexaFluor®488-labelled CREh was 5 nM in 10 mM MES pH 6.5, 150 mM NaCl, 10 mM $MgCl_2$, 0.05% Tween-20 (biophysical buffer) at 25 °C.

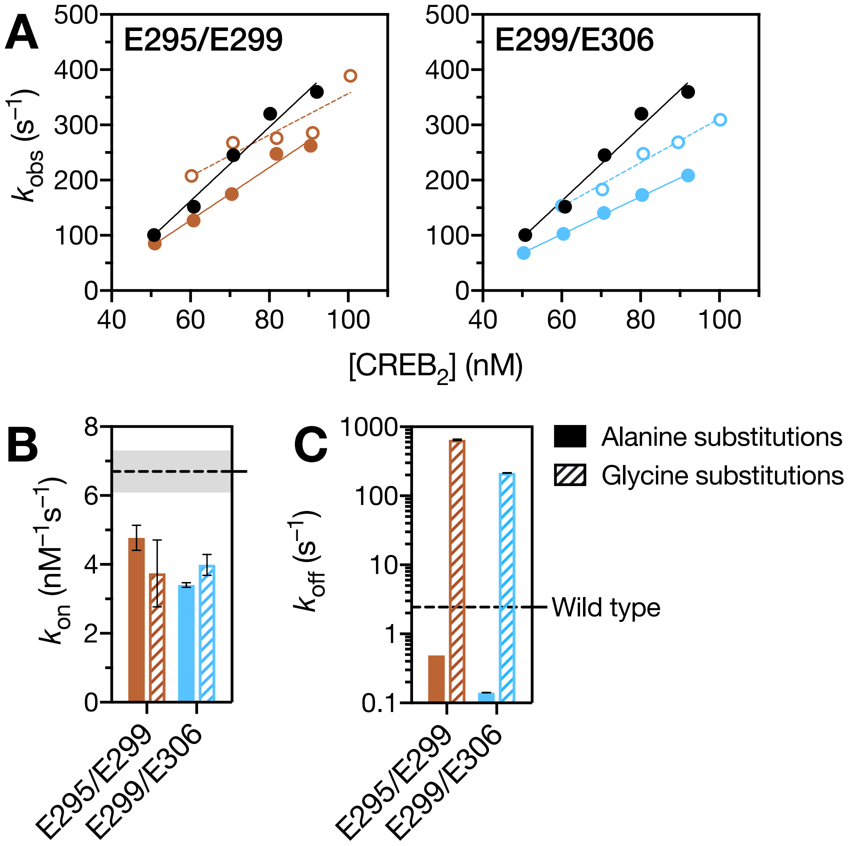


Supplementary Figure 9. Determination of kinetic rate constants for double mutants for double mutant cycle analysis. (A) Apparent association rate constants with 10 nM AlexaFluor®488-CREh for double mutants for residues E295/E299 and E299/E306. CREB $_{285}$bZIP is shown as a solid black circles, and double alanine mutants (coloured solid circles) and double glycine mutants (coloured open circles) for the two position pairs are shown in individual panels. Concentrations are for dimeric CREB. Lines are straight line fits and extracted rate constants (gradients) are shown in (B). (C) Dissociation rate constants for double mutants. Wild-type value is indicated by dotted line. Kinetic stopped-flow data were collected in 10 mM MES pH 6.5, 150 mM NaCl, 10 mM $MgCl_2$, 0.05% Tween-20 (biophysical buffer) at 25 °C.

**SUPPLEMENTARY METHODS**

**Derivation of equation for fitting homodimer refolding studies**

Let $P$ denote the protein monomer and $P_2$ – the protein dimer. The simplest chemical equation modelling their equilibrium is

$$2P \leftrightharpoons P_2. \qquad \text{(Eq. S1)}$$

Let $[P]$ and $[P_2]$ denote the concentrations of the respective species, and $P_T$ denote the total concentrations of protein. Then mass balance gives

$$P_T = [P] + 2[P_2]. \qquad \text{(Eq. S2)}$$

Let $L$ denote the equilibrium dissociation constant. When the system reaches equilibrium,

$$L = \frac{[P]^2}{[P_2]}. \qquad \text{(Eq. S3)}$$

Let $k_1$ and $k_2$ denote the association and dissociation rate constants, respectively, for homodimerization reaction Eq. S1. The rate of change in monomer concentration can be expressed as:

$$\frac{d[P]}{dt} = -k_1[P]^2 + 2k_2[P_2]. \qquad \text{(Eq. S4)}$$

At equilibrium $\frac{d[P]}{dt} = 0$, combining this with Eq. S3. shows the equilibrium constant $L$ is

$$L = 2\frac{k_2}{k_1} \qquad \text{(Eq. S5)}$$

and requiring mass balance (Eq. S2) leads to:

$$-k_1 dt = \frac{d[P]}{[P]^2 + \frac{L}{2}[P] - \frac{L}{2}P_T}. \qquad \text{(Eq. S6)}$$

Finding the roots of the quadratic expression in the denominator allows rewriting as:

$$-k_1 dt = \frac{d[P]}{\left([P] + \frac{L+2z}{4}\right)\left([P] + \frac{L-2z}{4}\right)}, \qquad \text{(Eq. S7)}$$

where

$$z = \sqrt{\frac{L^2}{4} + 2LP_T} \qquad \text{(Eq. 7a)}$$

Eq. S7. may then be integrated to obtain an expression for the concentration of monomeric protein with time:

$$[P] = \frac{\frac{L-2z}{4} - \left(\frac{L+2z}{4}\right) N e^{-k_1 z t}}{N e^{-k_1 z t} - 1}, \quad \text{(Eq. S8)}$$

where $N$ is a constant of integration that may be written in terms of the initial monomer concentration $[P]_0$ by evaluation of Eq. S8 at time zero:

$$N = \frac{\frac{L-2z}{4} + [P]_0}{\frac{L+2z}{4} + [P]_0}. \quad \text{(Eq. S8a)}$$

**Deviation of equation for equilibrium binding of a pre-formed protein dimer to DNA**

Let $D$ denote the DNA molecule and $P_2D$ – the protein-DNA complex. The binding of protein to DNA in our system can be described by the following set of equations:

$$2P \leftrightharpoons P_2, \quad \text{(Eq. S9)}$$
$$P_2 + D \leftrightharpoons P_2 D. \quad \text{(Eq. S10)}$$

Let $[D]$ and $[P_2D]$ denote the concentrations of the respective species, and $D_T$ denote the total concentrations of protein and DNA. The mass balance gives

$$P_T = [P] + 2[P_2] + 2[P_2 D], \quad \text{(Eq. S11)}$$
$$D_T = [D] + [P_2 D]. \quad \text{(Eq. S12)}$$

Let $K$ denote dissociation constants of the protein-DNA complex and $L$ denote dissociation constant of the protein dimer. When the system reaches equilibrium,

$$L = \frac{[P]^2}{[P_2]}, \quad \text{(Eq. S13)}$$
$$K = \frac{[P_2][D]}{[P_2 D]}. \quad \text{(Eq. S14)}$$

Substitution of Eqs. S12-S14 into Eq. S21 yields an expression for $[P_2D]$:

$$[P_2 D] = \frac{1}{2} P_T - \frac{1}{2} \sqrt{L \frac{K_1 [P_2 D]}{D_T - [P_2 D]}} - \frac{K_1 [P_2 D]}{D_T - [P_2 D]}. \quad \text{(Eq. S15)}$$

The anisotropy of a mixture $r$ is the sum of individual anisotropies of emitting species $r_i$ weighted by their fractional intensities $f_i$:

$$r = \sum r_i f_i = r_D f_D + r_{P_2 D} f_{P_2 D}, \quad \text{(Eq. S16)}$$

where

$$f_D + f_{P_2 D} = 1. \quad \text{(Eq. S16a)}$$

Eqs. S26 and S27 can be combined and rearranged as follows:

$$r = r_D + \Delta r f_{P_2 D}, \quad \text{(Eq. S17)}$$

where
$\Delta r = r_{P_2D} - r_D$. (Eq. S17a)

If binding of protein to DNA does not affect its fluorescence intensity, the fractional intensities can be re-written as molar fractions:

$$r = r + \frac{\Delta r[P_2D]}{D_T}. \quad \text{(Eq. S18)}$$

Otherwise, the anisotropy can be adjusted as described in reference[*]

Substituting Eq. S15 into Eq. S18 gives

$$r = r_D + \frac{\Delta r}{D_T}\left(\frac{1}{2}P_T - \frac{1}{2}\sqrt{L\frac{K_1[P_2D]}{D_T-[P_2D]}} - \frac{K_1[P_2D]}{D_T-[P_2D]}\right). \quad \text{(Eq. S19)}$$

Solving Eq. 18 for [$P_2D$] and substituting into Eq. S19 yields the final formula:

$$r = r_D + \frac{\Delta r}{D_T}\left(\frac{1}{2}P_T - \frac{1}{2}\sqrt{L\frac{Kz}{D_T-z}} - \frac{Kz}{D_T-z}\right), \quad \text{(Eq. S20)}$$

where
$$z = \frac{D_T(r-r_D)}{\Delta r}. \quad \text{(Eq. S20a)}$$

Eq. S20 is defined in an implicit form as $r = f(P_T, r)$, and the solution for $r$ was obtained using numerical methods.

* Crabtree MD and Shammas SL, "Stopped-Flow Kinetic Techniques for Studying Binding Reactions of Intrinsically Disordered Proteins", Methods Enzymol. **2018** *61*, 423-457
